# eCRUIS captures RNA-protein interaction in vitro and in vivo

**DOI:** 10.1101/2022.10.05.510920

**Authors:** Ziheng Zhang, Yuanbing Zhang, Ji-Long Liu

## Abstract

As an information bridge between DNA and protein, RNA regulates cellular processes and gene expression in a variety of ways. From synthesis to degradation, RNA is associated with a series of RNA-binding proteins. Therefore, it is very important to develop innovative methods to study the interaction between RNA and protein. Previously, we developed an RNA-centric method, called <u>C</u>RISPR-based <u>R</u>NA-<u>U</u>nited <u>I</u>nteracting <u>S</u>ystem (CRUIS), to capture RNA-protein interaction in cells. On this basis, here we develop an enhanced CRUIS (eCRUIS) by combining the power of dCas13d and the engineered promiscuous ligase TurboID. The new version allows us to label RNA-binding proteins on the target RNA within 30 minutes, which may be used in vivo. By introducing bait-assay with exogenous RNA, we confirm that eCRUIS can effectively label RNA-binding proteins on bait RNA in a short time. eCRUIS provides a wider range of in vitro and in vivo applications.

## INTRODUCTION

When the messenger transmits gene information from DNA to protein, the abundance, stability, and behavior of RNA are strictly regulated. As important RNA regulators, RNA-binding proteins have attracted much attention [1-4]. Strategies for studying RNA protein interaction can be roughly divided into two categories: protein-centric methods and RNA-centric methods [5, 6].

The protein-centric methods widely used in the past decades mainly include the well-known RIP (RNA-immunoprecipitation), CLIP (Cross-linking and immunoprecipitation), and their extended approaches. These approaches rely on the specific antibody of proteins of interest, capture and purify RNAs interacting with the target protein through immunoprecipitation of the RNA-proteins complex, and obtain the information about the type and abundance of these binding RNAs through RNA sequencing [7-9].

The RNA-centric methods study which proteins interact with RNA of interest. The development of this kind of method started late, but it has received extensive attention in recent years. The main research strategies include tracking and capturing target RNAs through RNA aptamers and antisense oligonucleotide probes [6].

For RNA-aptamer strategies, these aptamer sequences usually have specific secondary structures, can be recognized and bound by specific binding proteins or ligands, and have strong binding ability and high specificity. By inserting the RNA-aptamer sequence into the RNA of interest, the target RNA can be recognized by the corresponding binding protein. Some RNA-aptamers have been successfully applied to study the interaction between RNA and protein. For example, the MS2-MCP pair and the λ-N-BoxB pair have been used to identify and capture target RNAs [10, 11].

Another RNA centric strategy for studying RNA-protein interactions is based on antisense oligonucleotide probes. This type of method requires the design of a large number of DNA probes for the target RNA. Biotin-labeled DNA probes are usually synthesized to facilitate the subsequent enrichment of the target RNA-protein complexes through affinity streptavidin magnetic beads [12, 13]. Because probes binds to target RNA through base complementary pairing, their binding ability is much weaker than that between RNA-aptamer and ligands, so it is usually necessary to design a large number of probes to improve the efficiency of target RNA capture.

In addition to these two strategies, with the discovery of RNA targeting Cas nucleases, a class of CRISPR based methods for studying RNA-protein interactions was developed. We have previously developed the <u>C</u>RISPR-based <u>R</u>NA-<u>U</u>nited <u>I</u>nteracting <u>S</u>ystem (CRUIS), which the RNA-targeted Cas nuclease dCas13a and the proximity labeling system PUP-IT [14]. dCas13a, which losses of RNA cutting activity, acts as an RNA targeting element for recognizing and binding target RNA. The proximity labeling enzyme PafA is fused with dCas13a, and PupE is used to covalently label surrounding proteins.

Subsequently, a series of similar methods were reported. CRUIS and similar methods are based on RNA targeting Cas nucleases and proximity labeling systems to study the interaction between RNA and protein [14-16]. These methods make use of the ability of the Cas13 family to bind to target RNAs, and can efficiently recognize and capture target RNAs through the proximity labeling enzymes fused to it, and label the interacting proteins in living cells under physiological conditions.

In this study, we develop an enhanced CRUIS (eCRUIS) to capture the interaction between RNA and protein. eCRUIS can be applied not only in culture cells, but also in living organisms.

## RESULTS

### The strategy of eCRUIS

In order to quickly capture the RNA protein interaction of target RNA in living cells, our strategy is to replace the proximity labeling element of CRUIS with a system that can label proximity proteins in a short time. A recent study reported an engineered biotin ligase proximity labeling system, TurboID, which is based on the directed evolution of BioID [17]. TurboID exhibits extremely strong proximity labeling activity, which can effectively label surrounding proteins in less than 30 minutes, and is widely used to study protein-protein interactions in vivo in various species [17-20].

At the same time, with the further exploration of Cas13 family members, a series of CRISPR members have been discovered recently [21-24]. Among them, Cas13d shows a strong ability to cleave target RNA. Because the binding with the target RNA is the prerequisite for cleaving the target RNA, Cas13d is considered to have extremely high RNA binding activity.

Therefore, here, we developed an iterative version of CRUIS by combining dCas13d and TurboID, and named it eCRUIS. The use of TurboID in eCRUIS enables it to label adjacent proteins in a short period of time due to its proximity tagging element.

### eCRUIS implements short-term labeling

To test its proximity-tagging activity, we first constructed a stable cell line with Dox-inducible expression driven by the TRE promoter. After adding Dox to induce eCRUIS expression for 24 hours, biotin was added to promote the proximity labeling of surrounding proteins, and the efficiency of biotin labeling was detected by Western blotting (**Figure 1A**). Not surprisingly, within 30 minutes after the addition of exogenous biotin, it showed a strong biotin labeling signal. Without the addition of extra biotin, it showed a low labeling background (**Figure 1B**).

**Figure 1.**
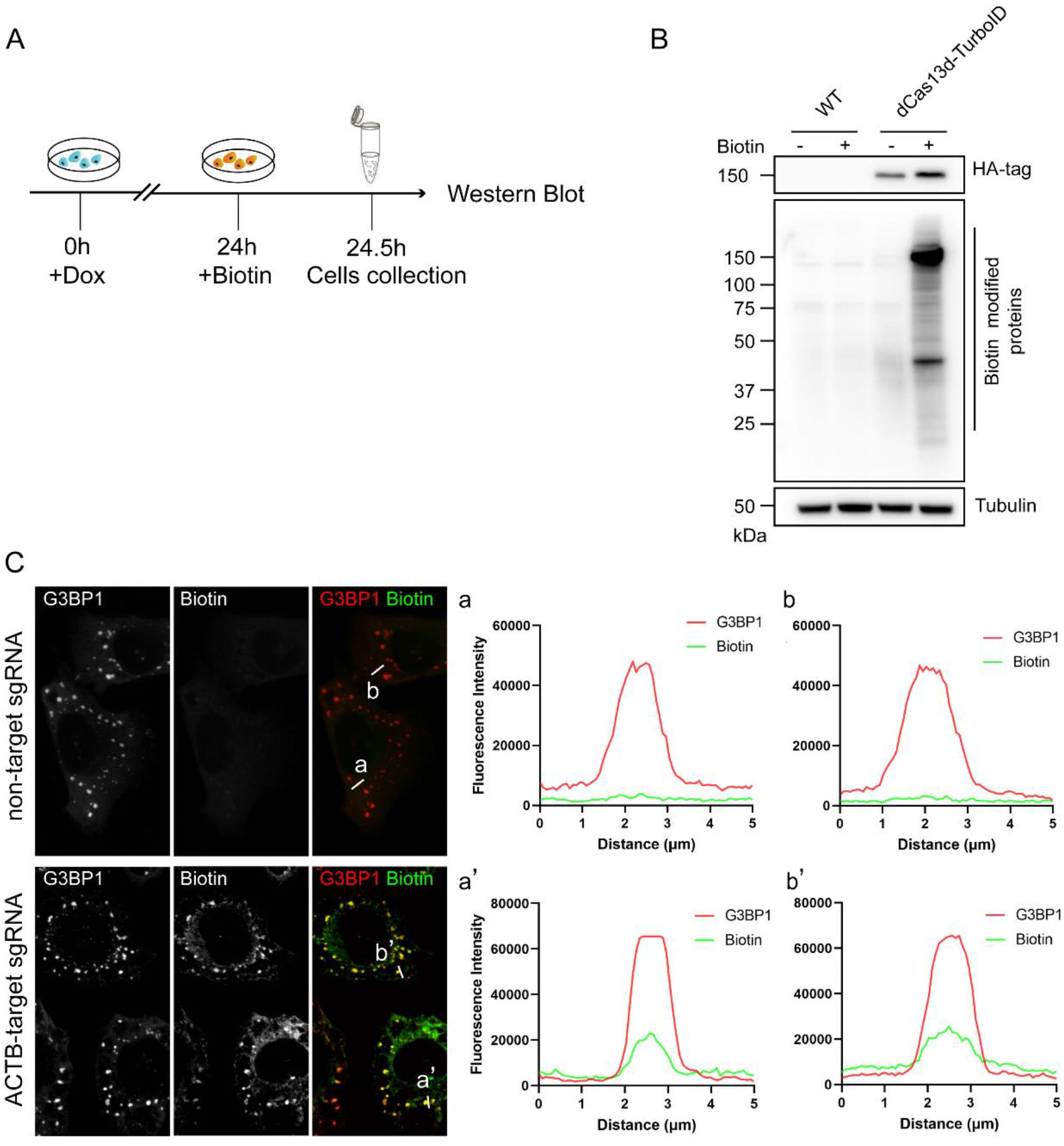
Construction of enhanced CRUIS (eCRUIS). (A) Timeline for testing eCRUIS proximity labeling activity. (B) proximity labeling activity testing for eCRUIS. (C) Tracking and labeling stress granules via eCRUIS. (A) Non-target sgRNA cannot guide eCRUIS to track and label stress granules. The sgRNA targeted to *ACTB* can guide eCRUIS to track the stress granules and label the stress granules with biotin within 30 min.

### eCRUIS binds to target RNA effectively

Next, we tested whether eCRUIS could bind to the target RNA under the guidance of specific sgRNA, and label the proteins around the target RNA through the stress granules tracking and labeling assay. We first constructed a stable cell line expressing eCRUIS in U2OS cells, and transfected the expression vector of ACTB-target sgRNA into the cells. After 24 hours of transfection, sodium arsenite induced the formation of stress granules. At the same time, we added biotin to the medium to promote the proximity-labeling of proteins.

The results showed that G3BP1, as a marker of stress granules, had good co-localization with biotin signal (**Figure 1C**). This shows that eCRUIS can effectively bind to target RNA and biotinylate surrounding proteins. It also indicates that the supplied biotin can effectively facilitate the proximity-labeling of eCRUIS and the background of biotin signal.

### eCRUIS captures RNA-interacting proteins in a short time

In order to further investigate whether eCRUIS can effectively capture proteins interacting with target RNAs, we designed a bait assay. First, we artificially designed a bait RNA containing multiple MS2 stem loops and expressed it in cells. At the same time, we expressed the MCP-BFP-V5 fusion protein into cells. Because of the natural interaction between the MS2 stem-loop and MCP, MCP-BFP-V5 will bind to the MS2 stem-loop on the artificially designed bait RNA.

We designed a vector to express bait RNA and MCP-BFP-V5. The design of bait RNA was based on the previously reported ultra-stable MS2 stem-loop scaffold [25]. The secondary structure of bait RNA was predicted by RNAFold, and the results showed that the MS2 stem-loop could be stably formed (**Figure 2A**). After testing, bait RNA could be efficiently expressed in cells (**Figure 2B**).

**Figure 2.**
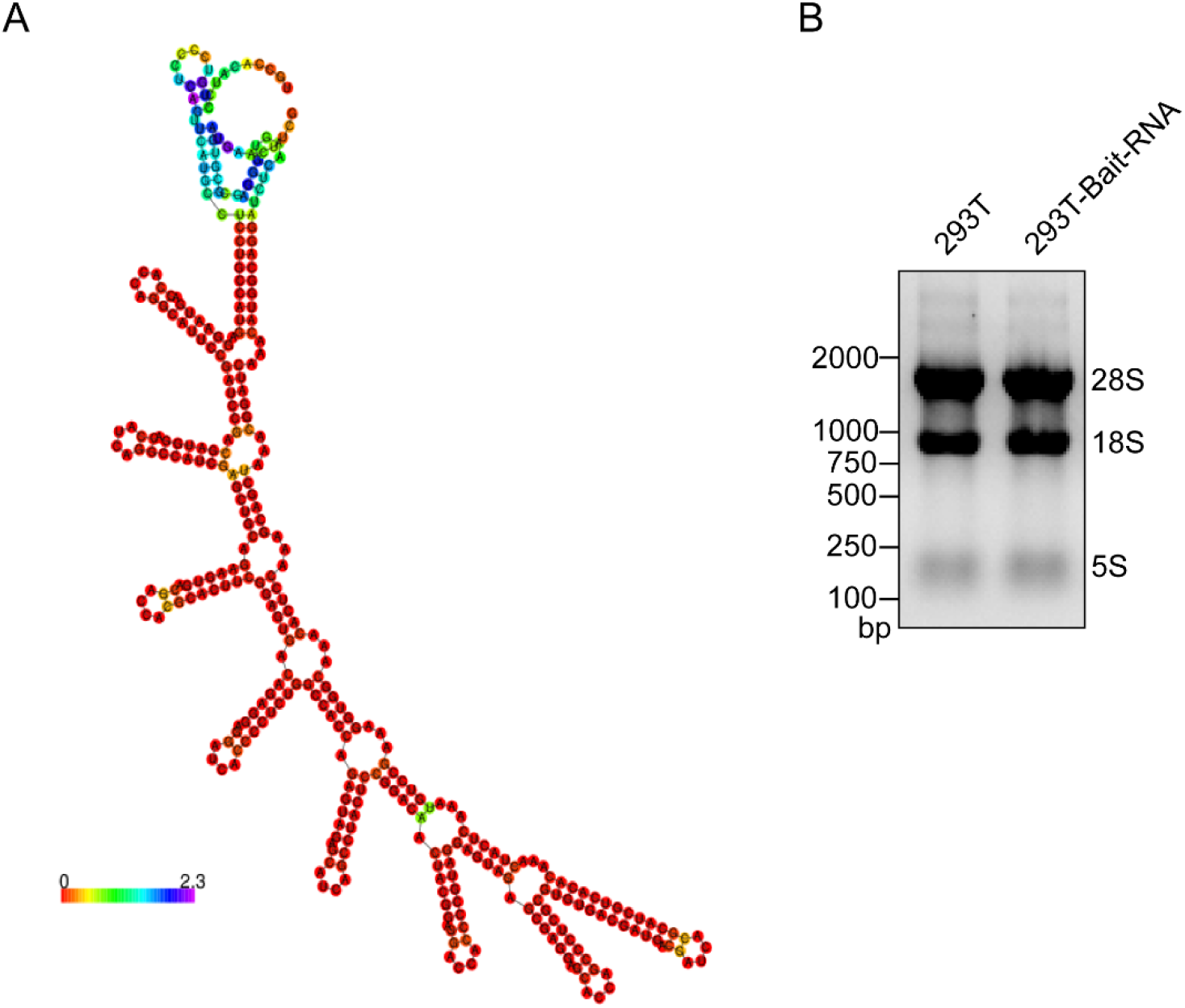
(A) Minimum free energy secondary structure of the Bait-RNA predicted by RNAflod. (B) Quality control by RNA electrophoresis of the total RNA.

At this time, under the guidance of bait RNA-specific sgRNA, eCRUIS can effectively label the MCP-BFP-V5 fusion proteins that binds to the MS2 stem-loop (**Figure 3A**). The sgRNA was driven by the human U6 promoter (**Figure 3B**). We first tested the expression of bait RNA in cells to determine that it could work normally and efficiently in cells. Bait RNA showed excellent expression levels 24 h after transfection (**Figure 3C**).

**Figure 3.**
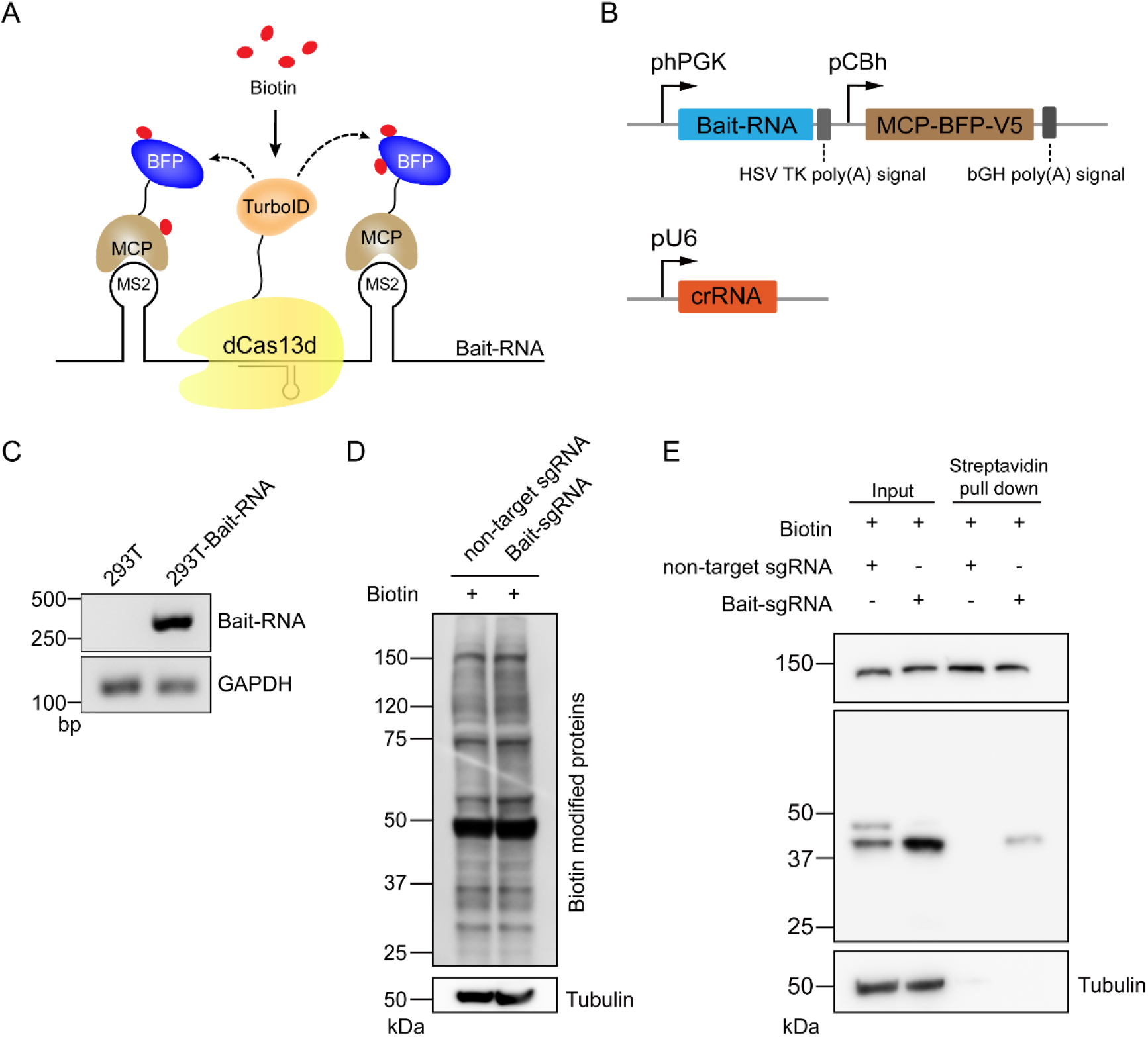
Bait assay verify eCRUIS can label interacting proteins of specific RNA within short time. (A) Design of Bait assay, express bait-RNA containing MS2 stem loop, and MCP-BFP fusion protein at the same time in cells. the sgRNA targeted to the bait RNA can guide the eCRUIS to label the MCP-BFP fusion protein. (B). Expression vectors used in bait assay. (C) Expression ability testing of Bait RNA. (D) Overall biotin signal. (E) sgRNA target to Bait-RNA could make eCRUIS label MCP-BFP efficiently.

Then, we conducted a decoy experiment. This was only to guide eCRUIS to use sgRNA specific to bait RNA to bind to bait RNA. Then, the MCP-BFP-V5 fusion protein bound to the bait RNA was labeled. Non-target sgRNA served as a negative control here, and finally enriched and detected by streptavidin beads.

After 36 h of transfection with bait-RNA and sgRNA, we added biotin to cells to promote the proximity labeling activity of TurboID, and ended labeling and collected samples 30 min after biotin addition. From the experimental results, eCRUIS exhibited obviously activity in all biotin labeling within 30 min (**Figure 3D**). MCP-BFP-V5 could be effectively labeled by biotin in the experimental group of Bait-sgRNA, while no obvious biotinylated MCP-BFP-V5 was observed in the non-target sgRNA control group (**Figure 3E**). These results suggest that eCRUIS can effectively label target RNA interacting proteins in a short time (30 min), and addind additional biotin can effectively control the background of random labeling.

### eCRUIS captures RNA-interacting proteins in vitro

Next we tested proximity labeling of eCRUIS in *Drosophila* S2 cells. We initially overexpressed dCas13d-TurboID-V5 recombinant protein under Ac5.1 promoter in S2 cells, and wild-type S2 cells were used as the control group (**Figure 4**). No obvious signal was detected in the control group, but extensive and strong signals were observed in the dCas13d-TurboID containing exogenous biotin, while there was almost no biotin signal in the streptavidin-HRP blotting results (**Figure 4A**). The immunofluorescence results revealed that after staining with streptavidin-AlexaFluor488, the fluorescence signal of the control group was little, while that of the cells with exogenous biotin was stronger (**Figure 4B**).

**Figure 4.**
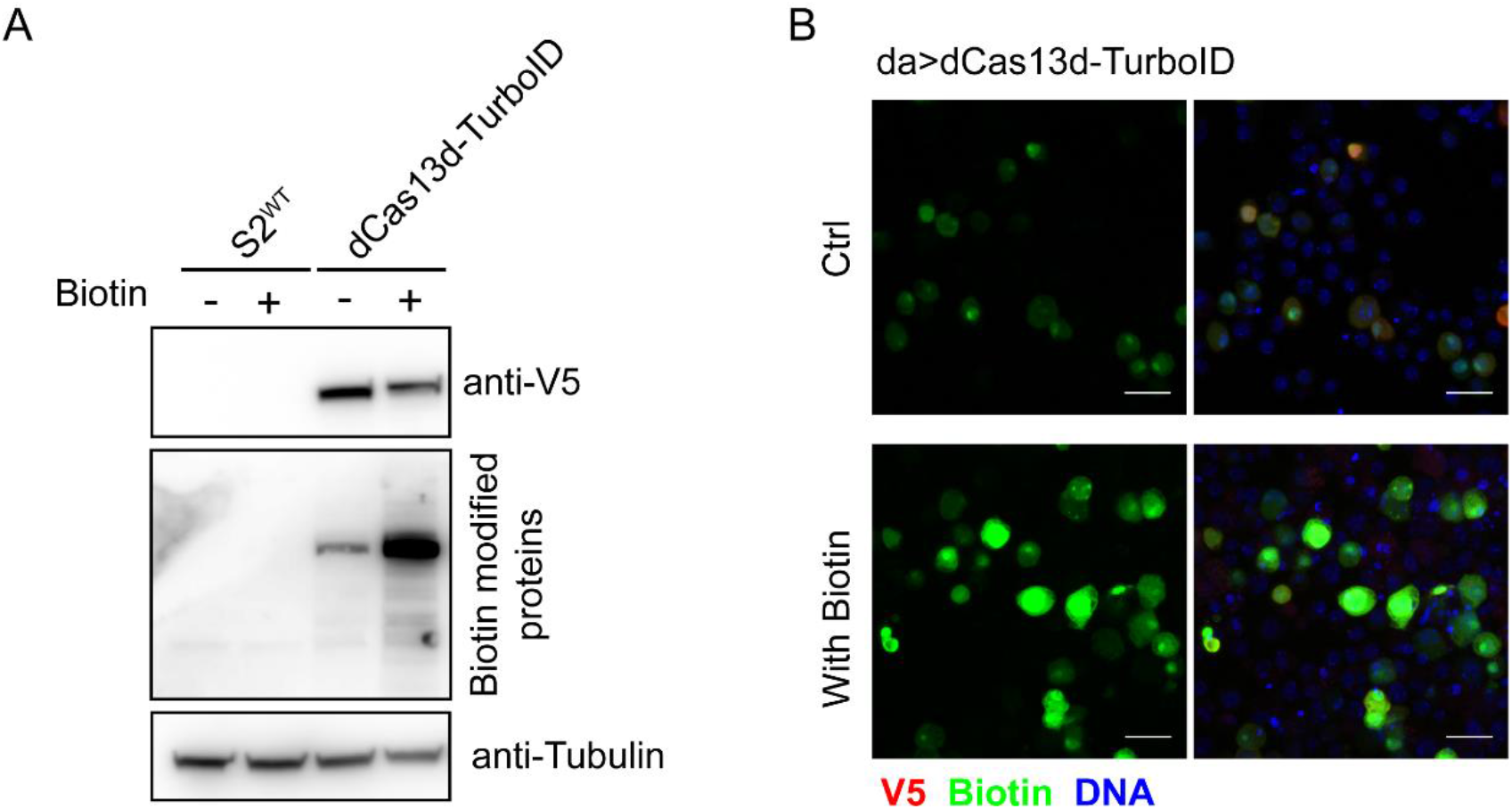
(A) Western Blot show that proximity labeling activity in S2 cells with or without biotin. (B) Immunofluorescence shows the proximity labeling activity in the S2 cells with or without biotin. Scale bars, 20 μm.

### eCRUIS captures RNA-interacting proteins in vivo

How to capture RNA-protein interactions in vivo has always been a key issue in understanding gene expression patterns. eCRUIS is easy to use and reusable, which is a natural advantage that can be applied in vivo. Therefore, we constructed eCRUIS transgenic *Drosophila* strains.

In order to verify the feasibility of the proximate labeling of eCRUIS in *Drosophila*, we used western blotting and immunofluorescence. We tried to generate transgenic flies and overexpressed dCas13d-TurboID-V5 recombinant protein ubiquitously under the drive of daughter-less Gal4. Here, flies raised in a normal diet served as the control group (**Figure 5A and B**). Streptavidin-HRP blotting results indicated that eCRUIS could biotinylate proteins with supplied biotin, because weak labeling signals could be detected in the control group, while signals from flies fed biotin were stronger (**Figure 5A**).

**Figure 5.**
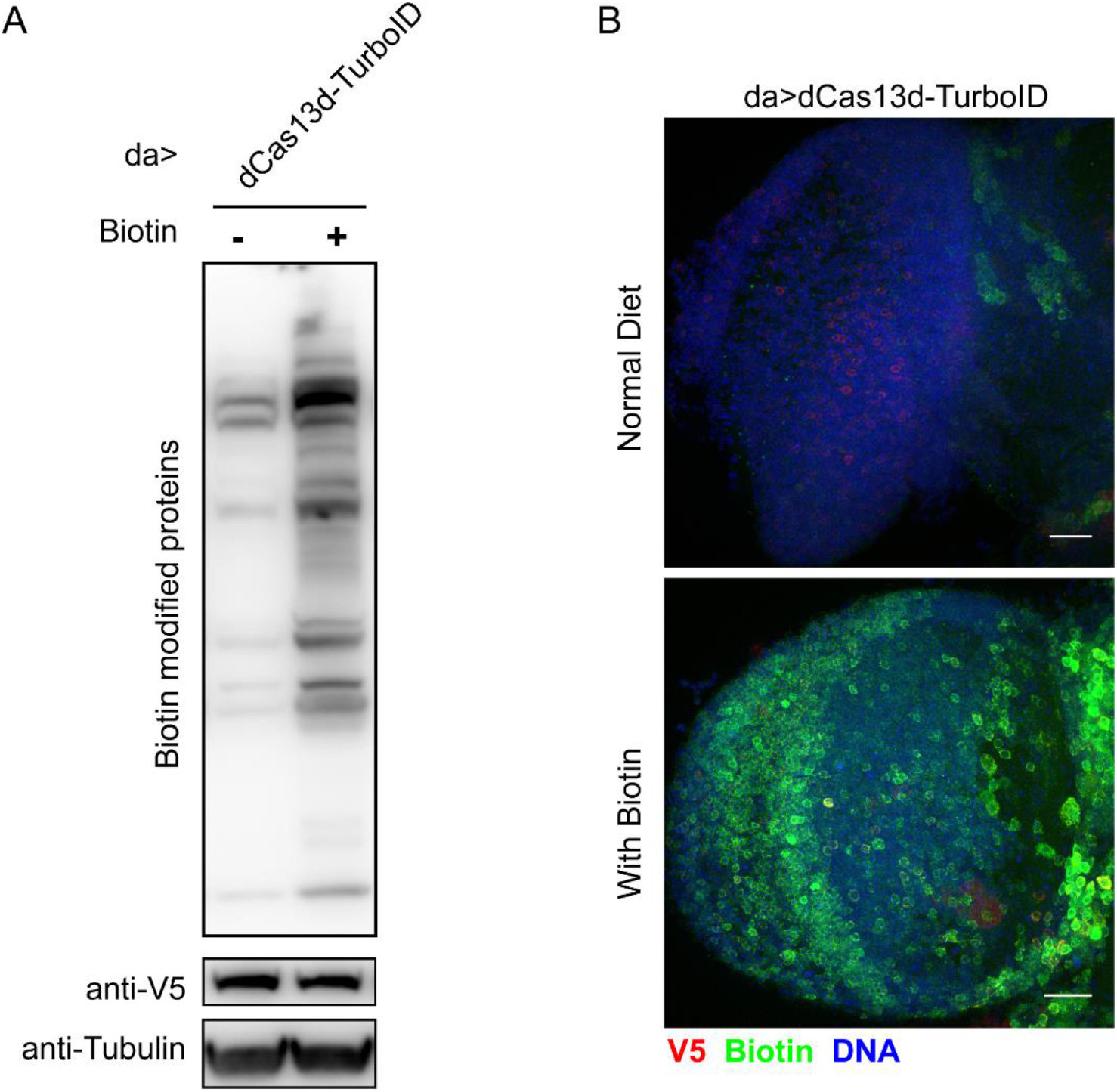
Canstrction of eCRUIS transgenic Drosophila and proximity labeling activity testing in vivo. (A) Western blotting shows that proximity labeling activity in vivo with or without feeding biotin. (B) Immunofluorescence shows the proximity labeling activity in the *Drosophila* brain with or without feeding biotin. Scale bars, 20 μm.

Then, we examined the wider application of eCRUIS in the *Drosophila* brain (**Figure 5B**). In the control group, few fluorescence signals were observed after staining with streptavidin-AlexaFluor488. In contrast, immunofluorescence images of flies fed biotin showed bright and extensive signals (**Figure 5B**). These results indicate eCRUIS can biotinylated proteins in situ and in vivo.

## DISCUSSION

As the intermediary between DNA and protein, RNA regulates gene expression and function in many ways. From biogenesis to degradation, RNA interacts with a series of RNA-binding proteins. This also suggests that the study of RNA function and its regulation mechanism must depend on the function and role of RNA-binding proteins in their entire life cycle [26, 27].

In the past decade, protein centered methods for studying RNA protein interactions have been widely developed, represented by RIP and CLIP. But this is not suitable for studying proteins that conversely interact with RNA of interest. Recently, RNA centered methods have been emphasized and developed. The main strategies include identifying and capturing target RNA and its interacting proteins through RNA aptamers and antisense oligonucleotide probes[6]. With the discovery of RNA targeting Cas nuclease, CRISPR based RNA protein interaction research methods have been developed.

The development of CRUIS and its iterative version is an effective supplement to the RNA centered method, which solves the shortcomings of the above two RNA centered methods (additional introduction of artificial engineering target RNA sequences and antisense oligonucleotides is required). Because of the natural advantages of Cas13 nuclease in binding to target RNA, it has realized the labeling and capture of RNA interacting with specific RNA in living cells.

Although CRUIS provides a novel strategy for capturing RNA-protein interactions, there is still room for optimization, such as the need for longer labeling time and the additional expression of PupE as a substrate for proximity labeling. In the initial version, we used PUP-IT as the adjacent tagging element, because PUP-IT needs to be linked to proximity proteins using PupE as a substrate. The Bio-PupE we used was achieved by fusing the BCCP domain to PupE [28]. It is auto-biotinylated. When the biotinylated sample is subsequently enriched by streptavidin beads, a large amount of free Bio-PupE unmarked on the target protein is enriched, resulting in excessive Bio-PupE in the mass spectrometry sample.

The quality of mass spectrometry data is affected by many invalid signals. The new version uses TurboID as a proximity labeling element. It uses biotin as a substrate to label surrounding proteins, and has a strong proximity labeling enzyme activity, which can quickly complete the labeling of adjacent proteins in less than 10 min [17]. In addition, before mass spectrometry sample preparation, cells are washed with PBS several times, which can effectively remove excess biotin. Even if free biotin binds to streptavidin-magnetic beads, it will not lead to the enrichment of invalid peptide signals during the preparation of mass spectrometry protein samples.

However, the existing methods and technologies are far from meeting the needs of researchers, especially in how to apply them to model organisms. Whether animals or plants, in different tissues and organs, with different developmental stages, different physiological or pathological states, RNA-protein interactions have huge differences, which is also the focus of research on RNA-protein interactions. It is technically difficult to obtain RNA-protein interactions in these states, and there is no effective method to follow.

The key to overcoming this technical challenge is to be able to apply methods to capture RNA-protein interactions in vivo. Compared with antisense oligonucleotide probes, eCRUIS has natural advantages in vivo application due to its bioencoding characteristics. As a promising method for in vivo use, eCRUIS has now successfully applied in model organisms.

CRUIS and its iterative version eCRUIS still have a lot of room for improvement, including how to reduce background, improve the labeling activity, and further improve specificity. The binding of Cas13 to the target RNA also forms a certain steric hindrance, which affects the binding of proteins interacting with this site. The existing methods still face various problems in research, and there is still a long way to go to study the interaction between RNA and protein.

## METHODS

### Plasmid construction and transgenic fly generation

The dCas13d and TurboID were cloned in the downstream of TRE3GV promoter in a lentivirus transfer vector by ClonExpress One Step Cloning kit (Vazyme). Here, an HA-tag was added in the 3’ end of TurboID, and (GGGGS)×2 is used as a linker between dCas13d and TurboID. P2A-EGFP was added in the C-terminal as the selection marker. For the plasmids for Drosophila, the Drosophila codon-optimized dCas13d and TurboID sequences containing V5 epitope tag were synthesized. Then digest pAc5.1 vector with EcoRI and NotI. The segments dCas13d, TurboID-V5 were inserted into the linearized pAc vector by seamless cloning according to ClonExpress One Step Cloning kit (Vazyme). The same approach can be used for the construction of pUAST dCas13d-TurboID-V5 for transgenic fly generation.

### Cell Culture and transfection

HEK293T cells and U2OS cells were grown in DMEM (Hyclone) supplemented with 10% FBS (Biological Industries) in a humidified incubator at 37°C with 5% CO_2_. Lipofectamine 3000 (Thermo) was used for transfection when the cell density is about 60% confluency, and all constructs used for transfection were prepared using E.Z.N.A.® Endo-free Plasmid DNA Mini Kit (Omega). For S2 cells, cells were cultured in Schneider’s Drosophila Medium containing 10% heat-inactivated FBS, 50 units penicillin G and 50 μg streptomycin sulfate per milliliter of medium in a 25°C sterile incubator. When the cells are approximately 80% confluent, change fresh media gently and transfect cells according to Effectene Transfection Reagent Kit (QIAGEN).

### Lentivirus production and infection

HEK293T cells was used for lentivirus production, psPAX2, pMD2.G and TetOn-eCRUIS-P2A-EGFP were transfected to HEK293T cells in the 3:1:4 ratio with Lipofectamine 3000 (Thermo), and change fresh medium gently after transfection 24 hours. Seventy-two hours after transfection, collect and filter the supernatant through a 0.45 μm PES filter before infection. For cell infection, U2OS cells and HEK293T cells were respectively plated in 60 mm dish, 24 hours after adding the supernatant with lentivirus, replace fresh medium with Dox (500ng/mL) to induce the expression of eCRUIS, and 48 hours after adding Dox, EGFP positive cells were collected by cell sorting.

### Western blotting

After treatment, 293T-TetOn-eCRUIS cells were washed with cold PBS and harvested by centrifuge. Lysis buffer (50 mM Tris 7.5, 150 mM NaCl, 1% Triton) with protease inhibitor was added to the pellet followed by incubating on ice for 30 min. Collect protein supernatant by centrifugation, followed by boiled with 1X protein loading buffer at 95°C for 10 min. Samples were separated by SDS-PAGE gels, followed by transferring proteins to PVDF membranes. After blocking with 5% BSA in TBST for 1 hour, the membrane was incubated with anti-HA antibody (Santa crue) and streptavidin-HRP (Cell Signaling) to identify the expression of dCas13d-TurboID fusion protein and the activity of TurboID.

For Drosophila, da gal4 flies were crossed with UAST dCas13d-TurboID-V5 flies and their embryos were raised at 25°C on either normal food or 100uM biotin-containing food accordingly. About 20 adult flies were collected and frozen by liquid nitrogen. Harvest cells by centrifugation after transfected 24-48 hours. Samples were lysed with 100uL RIPA buffer containing 1X protease inhibitor cocktail on ice for 1 hour. In Western blot, Streptavidin-HRP was used for detecting the activity of TurboID. Besides, anti-V5 antibody or anti-tubulin antibody and secondary antibodies correspondingly were used for verifying the expression of fusion protein and tubulin. All Western-blot signals were visualized via enhanced chemiluminescence and analyzed by Western Blot imaging system.

### Immunofluorescence

For stress granules tracking assay, after 24-hours transfection of sgRNA, 400 μM sodium arsenite and 400μM biotin were added to the medium. 30 min later, cells were washed by cold PBS 3 times, and fixed with 4% (w/v) paraformaldehyde in room temperature 15 min. After blocking with 5% BSA (diluted in 0.2% Triton X-100 in PBS) 1 hour in room temperature, anti-HA antibody (Santa crue) and streptavidin-Alexa Fluor 596 to identify the dCas13d-TurboId fusion protein and the biotin signal.

For *Drosophila*, about 10 brains from each group were dissected, then fixed with 4% (w/v) paraformaldehyde in PBS for 10min. For S2 cells, after 24 hours transfection, cells seeded on glass slides were fixed with 4% (w/v) paraformaldehyde in PBS for 10 min. Block samples with 5% bovine serum albumin and 0.2% Triton X-100 in PBS for 15 min. Samples were incubated with diluted Anti-V5 antibody in 0.2 % Triton X-100 in PBS and incubated overnight at room temperature. Then incubate samples with diluted Alexa Fluor 488-labeled secondary antibody, streptavidin-AlexaFluor488 and Hoechst 33342 in 0.2% Triton X-100 in PBS for about 4 h at room temperature. Transfer brains or glass slides to coverslips followed by adding ∼ 20 μL mounting medium and sealing slide with clear nail polish followed by confocal microscopy imaging.

## AUTHOR CONTRIBUTIONS

Z.Z. and J.L.L. conceived the studies. Z.Z. and Y.Z. performed the experiments. Z.Z. drafted the manuscript. J.L.L. revised the manuscript.

## ACKNOWLEDGEMENTS

We thank the Molecular Imaging Core Facility (MICF) at School of Life Science and Technology, ShanghaiTech University for providing technical support.

